# The GATA transcription factor Gaf1 represses tRNA genes, inhibits growth, and extends chronological lifespan downstream of fission yeast TORC1

**DOI:** 10.1101/700286

**Authors:** María Rodríguez-López, Suam Gonzalez, Olivia Hillson, Edward Tunnacliffe, Sandra Codlin, Victor A. Tallada, Jürg Bähler, Charalampos Rallis

**Author notes:** These authors contributed equally to this work (María Rodríguez-López, Suam Gonzalez). Correspondence: Charalampos Rallis; Jürg Bähler.

## Abstract

Target of Rapamycin Complex 1 (TORC1) signaling promotes growth and ageing. Inhibition of TORC1 leads to a down-regulation of factors that stimulate protein translation, including RNA polymerase III, which in turn contributes to longevity. TORC1-mediated post-transcriptional regulation of protein translation has been well studied, while analogous transcriptional regulation is less well understood. Here we screened fission yeast deletion mutants for resistance to Torin1, which inhibits TORC1 and cell growth. Mutants lacking the GATA transcription factor Gaf1 (*gaf1Δ*) grew normally even in high doses of Torin1. The *gaf1Δ* mutants shortened the chronological lifespan of non-dividing cells and diminished the lifespan extension triggered by Torin1 treatment. Expression profiling and genome-wide binding experiments showed that, after TORC1 inhibition, Gaf1 directly up-regulated genes for small-molecule metabolic pathways and indirectly repressed genes for protein translation. Surprisingly, Gaf1 bound to, and down-regulated the tRNA genes, so also functions as a transcription factor for genes transcribed by RNA polymerase III. We conclude that Gaf1 controls the transcription of both coding and tRNA genes to inhibit translation and growth downstream of TORC1.

## Introduction

The conserved Target of Rapamycin (TOR) signaling pathway is a key regulator for cellular growth and metabolism in response to nutrients and energy (Gonzalez and Rallis, 2017; González and Hall, 2017; Valvezan and Manning, 2019; Wei et al., 2013). In most organisms, TOR functions via two distinct multi-protein complexes, TORC1 and TORC2, which coordinate various aspects of growth and associated cellular processes (Hartmuth and Petersen, 2009; Ikai et al., 2011). TORC2 is not required for cell proliferation in fission yeast (*Schizosaccharomyces pombe*) but is involved in sexual differentiation, stress response, and actin function (Matsuo et al., 2007; Weisman and Choder, 2001). TORC1 activates protein synthesis and other anabolic processes and inhibits autophagy and other catabolic processes. Active TORC1 functions on lysosomes, or vacuoles in yeast, in response to growth or nutritional signals (Binda et al., 2009; Chia et al., 2017; Poüs and Codogno, 2011; Valbuena et al., 2012).

In all organisms tested, TORC1 signaling promotes ageing and shortens lifespan (Gonzalez and Rallis, 2017; González and Hall, 2017; Kaeberlein, 2010; Wei et al., 2013). Lifespan is influenced by multiple TORC1-dependent processes, including mitochondrial activity (Hill and Van Remmen, 2014), autophagy (Saxton and Sabatini, 2017), and protein translation (Bjedov and Partridge, 2011; Rallis et al., 2013). Global protein translation is controlled post-transcriptionally by TORC1, via direct phosphorylation of the ribosomal S6 kinases (S6K) and the translation factors eIF2α and 4E-BP (Ma and Blenis, 2009). Inhibition of S6K function can extend lifespan in fission yeast and other organisms (Bjedov et al., 2010; Rallis et al., 2014; Roux et al., 2006; Selman et al., 2009).

Besides post-transcriptional mechanisms, TORC1 promotes global translational capacity and ageing via transcriptional regulation (Valvezan and Manning, 2019). It stimulates the transcription of all ribosomal RNAs via both RNA polymerases I and III (Pol I and Pol III) (Iadevaia et al., 2014), although the mechanisms are poorly understood. Some evidence suggests that TORC1 regulates Pol I transcription via general transcription factors (Hannan et al., 2003; Mayer et al., 2004). TORC1 also regulates the conserved Maf1 protein which directly inhibits Pol III (Cai and Wei, 2015; Graczyk et al., 2018; Michels et al., 2010; Shor et al., 2010; Wei and Zheng, 2010; Wei et al., 2009). Pol III transcribes the highly abundant 5S ribosomal RNAs and transfer RNAs (tRNAs) which are central for translation, besides some other small RNAs (Arimbasseri and Maraia, 2016). Given the focus on protein-coding gene transcription, the regulation of Pol III transcription is less well understood. A recent study shows that Pol III activity limits lifespan downstream of TORC1 (Filer et al., 2017). Together, these findings suggest that TORC1-mediated control of transcription by Pol III is universally important for global translation and ageing. However, no specific transcription factors have been identified that bind to Pol III-dependent promoters and thus mediate translational control and lifespan.

Here we show that the conserved GATA transcription factor Gaf1 is required for TORC1-mediated suppression of cell growth in fission yeast. Upon TORC1 inhibition, Gaf1 not only binds to promoters of certain protein-coding genes, but also to the Pol III-transcribed tRNA genes which leads to their repression. Mutant cells lacking Gaf1 feature a shortened chronological lifespan. Our results uncover a transcription factor downstream of TORC1 that directly inhibits transcription of the tRNA genes, providing a global mechanism for transcriptional control of protein translation that prolongs lifespan.

## Results and Discussion

### Genes required for TOR-mediated growth inhibition

TORC1 and TORC2 can be selectively inhibited by Torin1, an ATP-analogue which blocks cell proliferation in fission yeast (Atkin et al., 2014; Thoreen et al., 2009). Using a low dose of Torin1 (5 μM), fission yeast mutants have recently been screened for resistance and sensitivity to reduced TOR signaling (Lie et al., 2018). Here we screened mutants under a four-fold higher dose of Torin1 (20 μM). This dose completely blocked cell growth (Fig. 1A) and reduced the size of both cells and vacuoles (Fig. 1B). Global protein translation was also reduced by Torin1, as reflected by reduced phosphorylation of ribosomal S6 protein and an increase in both total and phosphorylated forms of eIF2α (Fig. 1C). Together, these cellular phenotypes look like those triggered by combined caffeine and rapamycin treatment, which blocks TORC1 function (Rallis et al., 2013). We conclude that Torin1 leads to phenotypes that are diagnostic for strong TORC1 inhibition.

**Figure 1.**
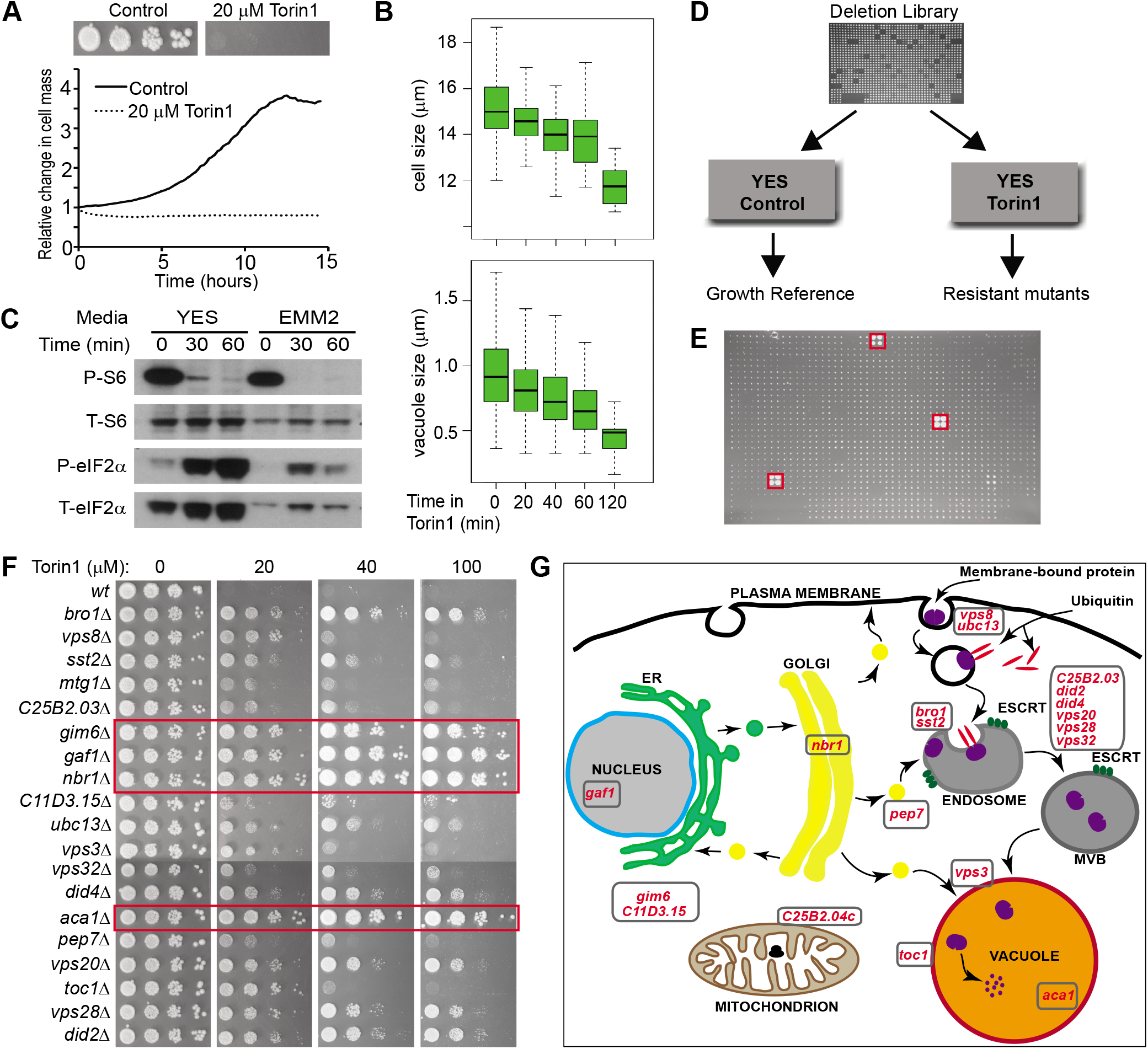
Genome-wide screen for Torin1-resistant mutants. **A.** Torin1 blocks cell proliferation. Top: ten-fold serial dilutions of wild-type cells spotted on rich solid medium; bottom: growth profiles in rich liquid medium using a microfermentor, in absence (Control) and presence of Torin1 as indicated. **B.** Torin1 leads to decreased cell and vacuole sizes. Cell size of septated wild-type cells (top) and vacuolar size (bottom) during 2hr treatments with Torin1. **C.** Torin1 alters phosphorylation status of translational regulators. Phosphorylated (P) and total amounts (T) of ribosomal S6 and eIF2α proteins in wild-type cells following 1hr Torin1 treatments in rich (YES) and minimal (EMM2) media as indicated. **D.** Design of genome-wide screens to identify mutants that are resistant to Torin1-mediated growth inhibition. We screened Bioneer versions 2 (3005 mutants) and 5 (3420 mutants) of genome-wide deletion libraries (Kim et al., 2010) in two independent repeats each, using 20 μM Torin1 on rich solid medium (YES). **E.** Representative example of deletion library plate with Torin1, containing 1536 colonies/plate with each mutant printed in quadruplicate. Red boxes indicate three Torin1-resistant mutants. **F.** Torin1 sensitivity test using spotting assays for a wild-type control (*wt*) and the 19 resistant mutants identified, applying increasing Torin1 concentrations as indicated. Red frames: 4 mutants showing the strongest resistance to all Torin1 concentrations tested. **G.** Scheme of cellular processes associated with the 19 genes identified that are required for Torin1-mediated growth inhibition (indicated in red).

We screened for deletion mutants that can suppress the strong growth inhibition by 20 μM Torin1. Two independent deletion libraries were screened in two repeats each (Fig. 1D). Fig. 1E shows examples of resistant mutant colonies. Overall, 19 mutants were resistant to Torin1-mediated growth inhibition in all 4 repeats (Table S1), 9 of which have been identified in the previous screen (Lie et al., 2018). We independently validated the 19 resistant mutant strains, both by PCR and backcrossing to a wild-type strain. The backcrossed mutants were spotted on plates with Torin1 to confirm that the drug-resistant phenotype was linked to the presence of the deletion cassette. While wild-type cells did not grow at all in Torin1, all 19 mutants managed to grow at least to some extend in the different concentrations of Torin1 (Fig. 1F). Four mutants were completely resistant to Torin1 at all concentrations, showing similar growth as on untreated medium (Fig. 1F, red frames).

Some mutants feature multi-drug resistance rather than resistance to specific drugs (Dawson et al., 2008). To exclude this possibility for the Torin1-resistant mutants, we assayed their growth in the presence of four other drugs (doxycycline, cadmium sulfate, bleomycin and cycloheximide). This analysis showed that all mutants were at least as sensitive to the other drugs as the wild-type control (Fig. S1A), indicating that their Torin1 resistance cannot be explained by multi-drug resistance. Could the resistance of some mutants simply reflect that they cannot take up Torin1? To exclude this possibility, we tested whether the Torin1-resistant mutants still showed any of the other phenotypes caused by TORC1 inhibition (Fig. 1B,C). The mutants still showed reduced levels of ribosomal S6 protein phosphorylation after Torin1 treatment, except *aca1Δ* (Fig. S1B), and decreased cell size, including *aca1Δ* (Fig. S1C). Together, these results indicate that Torin1 is taken up by the mutant cells which reveal different sensitivities to different TORC1 functions. Moreover, in all but the *aca1Δ* mutant, the growth resistance to Torin1 may be independent of translational control by ribosomal S6 phosphorylation.

The 19 genes identified in our screen function in a specific set of cellular processes (Fig. 1G; Table S1). Vesicular transport and vacuolar functions were associated with 13 genes, six of which encoding components of endosomal sorting complexes required for transport (ESCRT). Interestingly, many of these proteins are part of the recently discovered Nbr1-mediated vacuolar targeting (NVT) autophagic system (Liu et al., 2015). The NVT pathway does not contain core Atg proteins but depends on ESCRTs and the multi-vesicular body to deliver soluble cargoes to the vacuole. How might vesicular transport and the NVT pathway relate to TOR signaling? Disruption of the vesicle-mediated transport machinery at the endosome triggers a metabolic signature similar to TORC1 inhibition (Mulleder et al., 2016). It is possible that the NVT pathway is controlled by TORC1 or some of our resistant mutants affect the localization of TORC1 components to the vacuole, thus rendering the system more resistant to Torin1 inhibition. The remaining 6 genes identified in the screen encode proteins functioning in protein folding, mitochondrial ribosome assembly, the glutathione cycle, and arginine biosynthesis, as well as the TORC1-interacting protein Toc1p, and the GATA transcription factor Gaf1. In budding yeast, components of Golgi-to-vacuole trafficking are required for nitrogen- and TORC1-responsive regulation of GATA factors (Fayyadkazan et al., 2014; Puria et al., 2008). The screen results suggest that functional relationships between GATA factors and intracellular vesicles are conserved in fission yeast. Given our interest in TORC1-dependent gene regulation and the strong Torin1-resistance of *gaf1Δ* mutants (Fig. 1G), we further analyzed the function of Gaf1.

### Gaf1 is required for normal chronological lifespan and for full lifespan extension in Torin1-treated cells

TORC1 inhibition through nutrient limitation or rapamycin prolongs chronological lifespan in fission yeast (Rallis et al., 2013, 2014), defined as the time post-mitotic cells remain viable in stationary phase. Given that Gaf1 is required to arrest growth upon TOR inhibition, we hypothesized that Gaf1 may also play a role in chronological lifespan downstream of TORC1. To test this, we performed lifespan assays with *gaf1Δ* deletion mutant cells. Indeed, *gaf1Δ* cells were shorter-lived, with median and maximum lifespans of 3 and 16 days, respectively, compared to 5 and 20 days for wild-type cells (Fig. 2). Thus, Gaf1 is required for the normal lifespan of non-dividing cells.

**Figure 2.**
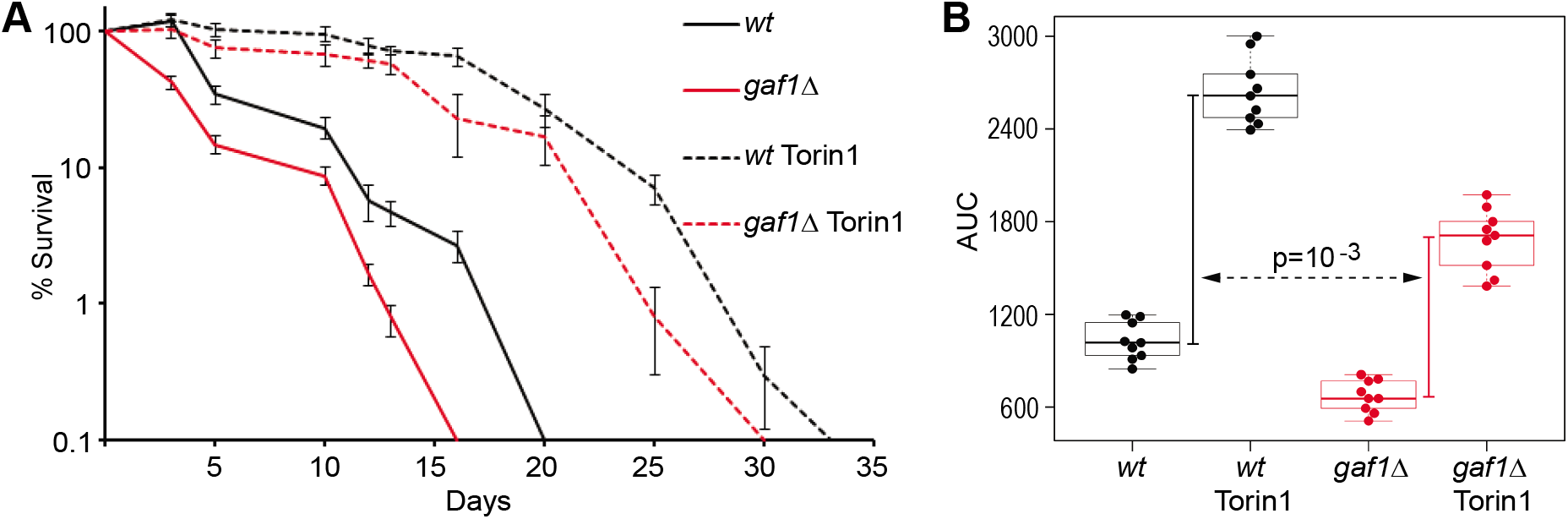
Gaf1 is required for normal chronological lifespan. **A.** Chronological lifespan assays in wild-type (*wt*) and *gaf1Δ* mutant cells grown in EMM2, grown in the absence or presence of 8 μM Torin1 as indicated. Error bars represent the standard deviation calculated from three independent cell cultures, with each culture measured three times at each timepoint. **B.** AUC for lifespan assays of *wt* and *gaf1Δ* mutant cells without or with Torin1 treatment as indicated. Vertical bars show the Torin1-mediated increase in average AUC values for *wt* (black) and *gaf1Δ* (red), with p value reflecting that the lifespan increase is significantly larger in *wt* than in *gaf1Δ* cells.

Torin1 increases lifespan in flies (Mason et al., 2018) and suppresses senescence in human tissue cultures (Leontieva and Blagosklonny, 2016). To analyze the effect of Torin1 on chronological lifespan in fission yeast, and any role of Gaf1 in this condition, we pre-treated exponentially growing wild-type and *gaf1Δ* cells with Torin1 and tested for subsequent effects on lifespan during stationary phase. Torin1 substantially prolonged lifespan in wild-type cells, with median and maximum lifespans of 18 and 33 days, respectively, compared to 5 and 20 days in untreated cells (Fig. 2A). In *gaf1Δ* cells, Torin1 also prolonged lifespan, but to a lesser extent than in wild-type cells, with median and maximum lifespans of ~13 and 30 days, respectively (Fig. 2A). To quantify the role of Gaf1 in Torin1-mediated longevity, we calculated the areas under the curve (AUC, measured as days × % survival) of the lifespan assays. In wild-type cells, the lifespan was prolonged from an average AUC of 1044 to 2689 (increase of 1645), whereas in *gaf1Δ* cells, the lifespan was prolonged to a lesser extent, from an average AUC of 681 to 1709 (increase of 1027) (Fig. 2B). We conclude that Gaf1 is also required for the full lifespan extension resulting from Torin1-mediated TOR inhibition during cell proliferation. However, Torin1 still can prolong lifespan to a considerable extent in the absence of Gaf1, indicating that other factors contribute to this longevity. Indeed, we have previously identified several proteins required for lifespan extension when TORC1 is inhibited, including the S6K homologue Sck2 (Rallis et al., 2014).

### Gaf1-dependent transcriptome regulation following TOR inhibition

Given that the Gaf1 transcription factor was essential for growth inhibition by Torin1 (Fig. 1G), we further analyzed its function in this condition. Gaf1 accumulated in the nucleus within a few minutes following Torin1 treatment (Fig. 3A). Consistently, Gaf1 is known to translocate to the nucleus during nitrogen limitation, which also inhibits TORC1 (Laor et al., 2015). This regulatory pattern for GATA transcription factors is conserved: budding yeast Gln3 and Gat1 (Broach, 2012) and mammalian GATA6 (Xie et al., 2015) are also sequestered in the cytoplasm by active TORC1 and translocate to the nucleus upon TORC1 inhibition.

**Figure 3.**
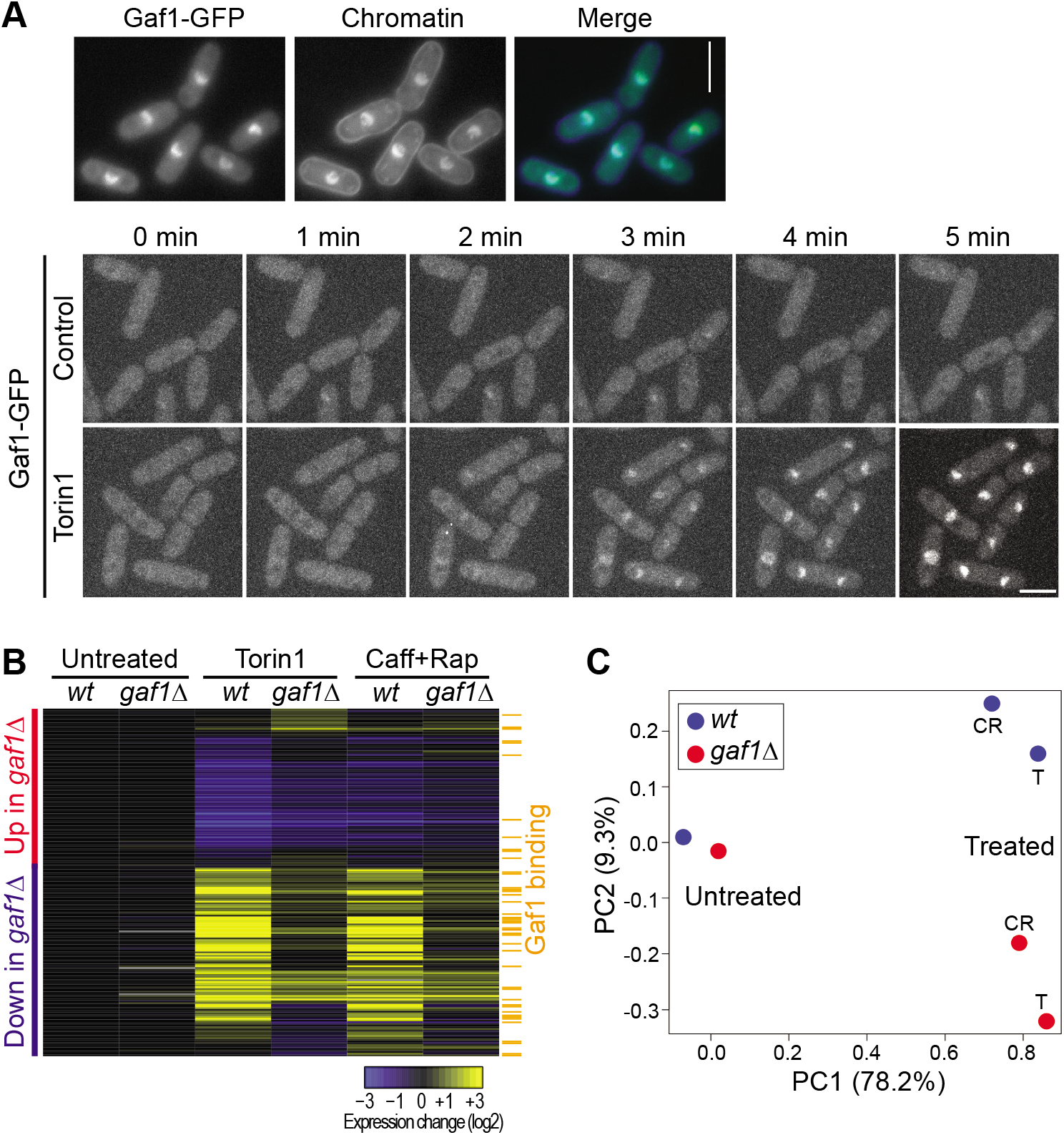
Gaf1-dependent gene expression. **A.** Top panels: Fluorescence microscopy of cells expressing GFP-tagged Gaf1 (left) with chromatin stained by Hoechst 33342 (middle) after 10 min exposure to 20 μM Torin1. Bottom panels: Fluorescence microscopy of live Gaf1-GFP cells, showing stack projections of 1-min time lapses in YES medium. Cells are shown before (0 min) and in 1-min intervals after addition of either DMSO (solvent control; upper panels) or Torin1 solution (20 μM final; lower panels). Gaf1-GFP is visible inside the nucleus in all cells within 3 min after Torin1 addition. Scale bars: 5 microns. **B.** Hierarchical clustering of microarray data. Columns represent wild-type (*wt*) or *gaf1* deletion mutants (*gaf1Δ*) before (Untreated) and after 1hr treatment with 20 μM Torin1 or with 10mM caffeine and 100 ng/ml rapamycin (Caff+Rap). Rows represent the 198 genes whose mRNA levels changed ≥1.5-fold in Torin1-treated *gaf1Δ* cells relative to wild-type cells, consisting of 90 genes showing higher expression in *gaf1Δ* cells (red bar) and 108 genes showing lower expression in *gaf1Δ* cells (blue bar). In untreated cells, only 3 genes showed ≥1.5-fold expression changes in *gaf1Δ* relative to wild-type cells. Average RNA expression changes (from 2 independent repeats) in the different genetic and pharmacological conditions relative to *wt* control cells are color-coded as shown in the legend at bottom. The orange bars at left indicate 43 genes whose promoters were bound by Gaf1 after 60 min with Torin1. **C.** Principal component (PC) analysis of all genes measured by microarrays. PC1 separates untreated cells from cells treated with Torin1 (T) or caffeine and rapamycin (CT), while PC2 separates wild-type (*wt*, blue) from *gaf1* deletion mutants (*gaf1Δ*, red). The percentages of the x- and y-axes show the contribution of the corresponding PC to the difference in the data.

In fission yeast, Gaf1 is known to activate genes functioning in amino-acid transport but represses *ste11*, encoding a master regulator for sexual differentiation (Kim et al., 2012; Ma et al., 2015). To systematically identify Gaf1-dependent transcripts before and after TOR inhibition, we performed microarray analyses of wild-type and *gaf1Δ* cells, both before and after Torin1 treatment. In the absence of Torin1, the expression signatures of wild-type and *gaf1Δ* cells were similar (Fig. 3B,C). We conclude that in proliferating cells Gaf1 plays no or a negligible role in gene regulation, consistent with the absence of Gaf1 nuclear localization when TORC1 is active (Fig. 3A; Laor et al., 2015).

Treatment with Torin1, on the other hand, resulted in substantial transcriptome changes in both wild-type and *gaf1Δ* cells (Fig. 3B,C). But in *gaf1Δ* cells, the expression signature triggered by Torin1 markedly differed from the signature in wild-type cells (Fig. 3B,C). Overall, 90 and 108 genes consistently showed ≥1.5-fold higher or lower expression, respectively, in *gaf1Δ* relative to wild-type cells after Torin1 treatment (Fig. 3B; Table S2). Cells treated with caffeine and rapamycin, which inhibit TORC1 but not TORC2 (Rallis et al., 2013), showed similar expression signatures as Torin1-treated cells, in both wild-type and *gaf1Δ* (Fig. 3B,C). This result indicates that the Torin1-mediated expression signatures in wild-type and *gaf1Δ* cells reflect TORC1 inhibition. We conclude that after TORC1 inhibition, Gaf1 affects the expression of ~200 genes, either positively or negatively.

We performed functional enrichment analyses for these Gaf1-dependent genes using AnGeLi (Bitton et al., 2015). The 90 genes that were higher expressed in *gaf1Δ* than in wild-type cells (i.e., genes repressed by Gaf1) were typically down-regulated in Torin1-treated wild-type cells, but less so in *gaf1Δ* cells (Fig. 3B). These genes were enriched in anabolic processes such as biosynthesis (61 genes, p=9.4×10^−10^), ribosome biogenesis (19 genes, p=1.6×10^−3^) and cytoplasmic translation (31 genes, p=1.0×10^−16^), including 25 genes encoding ribosomal proteins. Figure S2 provides a visualization of all Gene Ontology (GO) Biological Processes enriched among the 90 genes. Many of these genes are also repressed as part of the core environmental stress response (43 genes; p=1.4×10^−20^; Chen et al., 2003) and are highly expressed in proliferating cells (mean of 46.9 mRNA copies/cell *vs* 7.5 copies for all mRNAs, p=1,2×10^−26^; mean of 85,821 protein copies/cell *vs* 16,166 for all proteins, p=3.7×10^−19^; Marguerat et al., 2012).We conclude that upon TORC1 inhibition, Gaf1 contributes to the down-regulation of highly expressed genes functioning in global protein synthesis.

The 108 genes that were lower expressed in *gaf1Δ* than in wild-type cells (i.e., genes induced by Gaf1) were typically up-regulated in Torin1-treated wild-type cells, but less so in *gaf1Δ* cells (Fig. 3B). These genes were enriched in several metabolic/catabolic processes of small molecules, including organonitrogen compounds (43 genes, p=4.6×10^−14^) such as cellular amino acids (18 genes, p=4.1×10^−5^), urea (6 genes, p=7.3×10^−5^), and organic acids (20 genes, p=0.001). Figure S2 provides a visualization of all GO Biological Processes enriched among the 108 genes. There was also a substantial overlap with genes that are induced under nitrogen limitation (43 genes, p=1.3×10^−29^; Mata et al., 2002) and with genes that are periodically expressed during the cell cycle (41 genes, p=1.6×10^−12^, Marguerat et al., 2012), including 9 histone genes. These results suggest a Gaf1-dependent transcriptional program to adjust the metabolism of amino acids and other metabolites, possibly to recycle nutrients under conditions that do not allow rapid proliferation. Together, the expression profiling indicates that Gaf1 regulates physiological changes supporting the growth arrest triggered by TORC1 inhibition.

### Gaf1 binds to both coding and tRNA genes following TOR inhibition

The microarray analyses identified genes whose expression depends on Gaf1, some of which may be directly regulated by Gaf1. To detect genes whose promoters are bound by Gaf1, we performed chromatin immunoprecipitation followed by sequencing (ChIP-seq) of Gaf1-GFP cells, before and after Torin1 treatment. The number of gene promoters bound by Gaf1 increased from 165 before Torin1 treatment to 454 at 60 min after Torin1 treatment, with 93 genes in common between the two conditions (Fig. 4A). Gaf1 binding sites upstream of close, divergently expressed genes were assigned to both genes. The 454 Gaf1 target genes after Torin1 treatment consisted of 245 protein-coding genes and 209 non-coding genes. The full lists of genes whose promoters were bound by Gaf1 is provided in Table S3.

**Figure 4.**
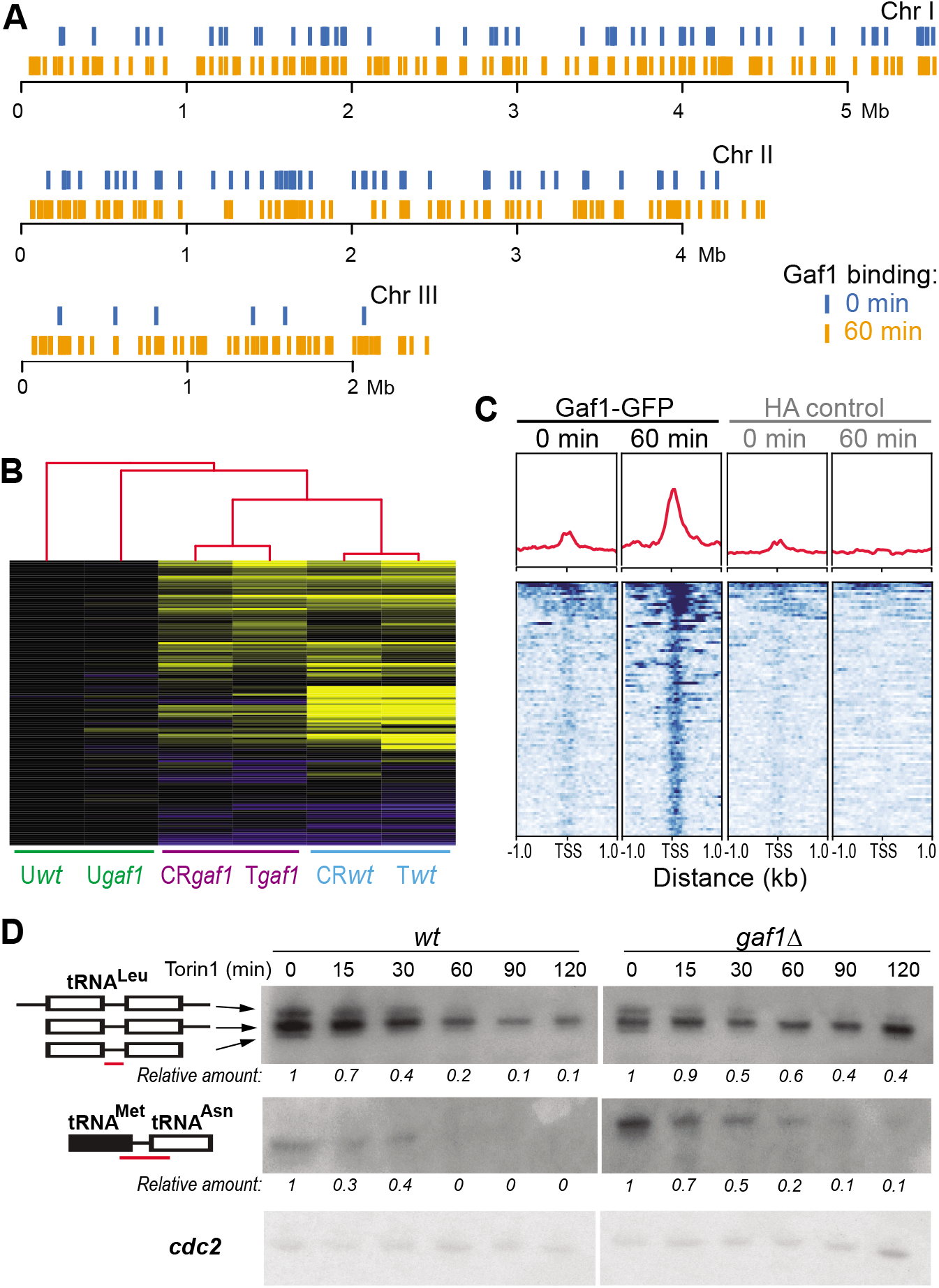
Gaf1 regulation of protein-coding and tRNA genes. **A.** Gaf1 binding sites across the three *S. pombe* chromosomes, before (0 min, blue) and after treatment with 20 μM Torin1 (60 min, orange). **B.** Hierarchical clustering of microarray expression data for 150 protein-coding genes bound by Gaf1 after Torin1 treatment and for which expression data were available for all conditions. The conditions have been clustered as well (red tree on top) and are grouped as follows: untreated wild-type and *gaf1Δ* cells (U*wt*, U*gaf1*), caffeine+rapamycin-or Torin1-treated *gaf1Δ* cells (CR*gaf1*, T*gaf1*), and caffeine+rapamycin- or Torin1-treated wild-type cells (*CRwt*, T*wt*). Expression changes are color-coded as described in Fig. 3B. **C.** Gaf1 shows increased binding to tRNA genes after Torin1 treatment. Top, red curves: average Gaf1-binding profiles aligned to the transcription start sites (TSS) of all *S. pombe* tRNA genes before (0 min) and after (60 min) Torin1 treatment as indicated, along with corresponding control ChIP-seq experiments (HA). Bottom: heatmaps of Gaf1 binding around the TSS of all 196 tRNAs, ordered by normalized ChIP-seq coverage. **D.** Northern blot analyses for leucine, methionine and asparagine precursor tRNAs from wild-type and *gaf1Δ* cells, treated with 20 μM Torin1 over 120 min as indicated. Probes to detect the precursor tRNAs are indicated in red at left (Otsubo et al., 2018). Probes for *cdc2* were used as a loading control. Relative tRNA amounts (normalized to time 0) have been measured using ImageJ.

AnGeLi analysis of the protein-coding target genes revealed significant enrichment in metabolic processes of organonitrogen compounds (55 genes, p=1.4×10^−6^), including nucleotides (24 genes, p=0.0009) and organic acids (34 genes, p=0.0003), among others. Figure S3 provides a visualization of all GO Biological Processes enriched among the 245 coding genes. The Gaf1 target genes were also enriched for genes induced under nitrogen limitation (40 genes, p=1.6×10^−11^) and periodically expressed during the cell cycle (53 genes, p=4.5×10^−6^). Overall, these target genes showed similar functional enrichments to the genes whose expression was induced by Gaf1. Accordingly, Gaf1 binding sites were enriched among the genes whose expression was induced by Gaf1 (Fig. 3B, orange bars). Moreover, most protein-coding genes bound by Gaf1 after Torin1 treatment were induced by Torin1 or by caffeine and rapamycin, but were less induced in *gaf1Δ* cells, leading to distinct clusters for wild-type and mutant conditions (Fig. 4B). We conclude that most coding Gaf1 target genes are transcriptionally up-regulated by Gaf1 upon inhibition of TORC1.

Notably, Gaf1 bound to promoters of 20 transcription factor genes (Table S3; Fig. S3). Typically, these factors were induced in wild-type cells after TORC1 inhibition, but less so in *gaf1Δ* cells. Many of these factors are involved in different stress responses or cell-cycle regulation, including Atf1 (Wilkinson et al., 1996), Cbf12 (Chen et al., 2003), Fep1 (Bekker et al., 1991), Fil1 (Duncan et al., 2018), Hsr1 (Chen et al., 2008), Klf1 (Shimanuki et al., 2013), Loz1 (Corkins et al., 2013), Pap1 (Chen et al., 2008), Php3 (Mercier et al., 2006) and Sep1 (Rustici et al., 2004). The large number of transcription-factor targets indicates that Gaf1 may indirectly control some Gaf1-dependent genes by regulating other transcription factors. Indeed, Gaf1 inhibited the expression of many genes functioning in translation (Fig. S2), but these genes were not among its direct binding targets. Thus, these genes may be indirectly regulated by Gaf1 via other transcription factors; for example, the Gaf1 target Atf1 is known to repress translation-related genes during stress (Chen et al., 2008), raising the possibility that it also represses these genes during TORC1 inhibition in a Gaf1-dependent manner.

The 209 non-coding genes among the Gaf1 target genes included 82 tRNA genes and a snoRNA involved in tRNA regulation, besides large non-coding RNAs (Table S3). In fact, coverage plotting indicated that Gaf1 binds to all tRNA genes, which are clustered in fission yeast (Fig. 4C; Fig. S4). The binding was enriched over the transcription start sites for tRNA genes and strongly increased after Torin1 treatment (Fig. 4C). We conclude that Gaf1 binds not only to Pol II transcribed genes, but also to tRNA genes that are transcribed by Pol III. Does Gaf1 repress or activate the tRNA genes? To address this question, we performed northern analyses to determine the expression of three tRNA genes as a function of Gaf1 and Torin1. The abundant mature tRNAs are rapidly processed from precursor tRNAs, so any expression changes of tRNAs requires detection of its precursors (Otsubo et al., 2018). The expression of tRNA precursors decreased during Torin1 treatment in wild-type cells, while in *gaf1Δ* cells this expression decrease was delayed and less pronounced (Fig. 4D). We conclude that upon TOR inhibition Gaf1 binds to tRNAs and inhibits their expression.

Down-regulation of precursor tRNA expression is required for TORC1 inhibition during nitrogen starvation in fission yeast (Otsubo et al., 2018), indicating that tRNAs can act upstream of TORC1. Our experiments, on the other hand, point to a mechanism of tRNA regulation downstream of TORC1. Together, these findings indicate a regulatory feedback mechanism, involving precursor tRNAs, TORC1 and Gaf1, to ensure that tRNA expression matches the physiological requirements. Our results reveal a transcription factor that globally inhibits Pol III-mediated expression of tRNAs, along with Pol II-mediated expression of protein-coding genes functioning in translation- and metabolism-related processes. It will be interesting to test whether this function is conserved for the orthologous GATA transcription factors.

## Conclusion

The GATA transcription factor Gaf1 is essential for the block of cell proliferation triggered by Torin1; in its absence cell growth is not reduced, even in high doses of Torin1 (Fig. 1G). Gaf1 is also required for normal chronological longevity, and it may contribute to, but is not absolutely necessary for the extended lifespan we observed in Torin1-treated cells (Fig. 2). Upon TORC1 inhibition, Gaf1 inhibits the expression of diverse genes functioning in protein translation, including protein-coding genes which seem to be indirectly controlled by Gaf1 as well as tRNA genes which are binding targets of Gaf1 (Figs. 3 and 4). Gaf1 also positively controls genes functioning in metabolic pathways for nitrogen-containing molecules, which support the physiological adaptation to lowered protein synthesis. Thus, Gaf1 can directly regulate both Pol II- and Pol III-transcribed genes. It is possible that Gaf1 elicits its repressor activity at tRNA genes by recruiting a histone deacetylase: previous work in fission yeast has identified potential loading sites for components of a Clr6 histone deacetylase complex at tRNAs (Zilio et al., 2014). Down-regulation of global protein translation is beneficial for longevity in all organisms studied, including fission yeast (Kaeberlein and Kennedy, 2011; Rallis and Bähler, 2013). Given the role played by Gaf1 in inhibiting translation-related factors, Gaf1 may inhibit ageing by contributing to the down-regulation of translation upon TORC1 inhibition (Fig. 5). Gaf1 thus defines a parallel, transcription-based branch of translational and metabolic control downstream of TORC1, in parallel to the post-translational branch exerted by translational regulators like S6K and others (Fig. 5). This branch is essential for the growth inhibition triggered by lowered TORC1 activity.

**Figure 5.**
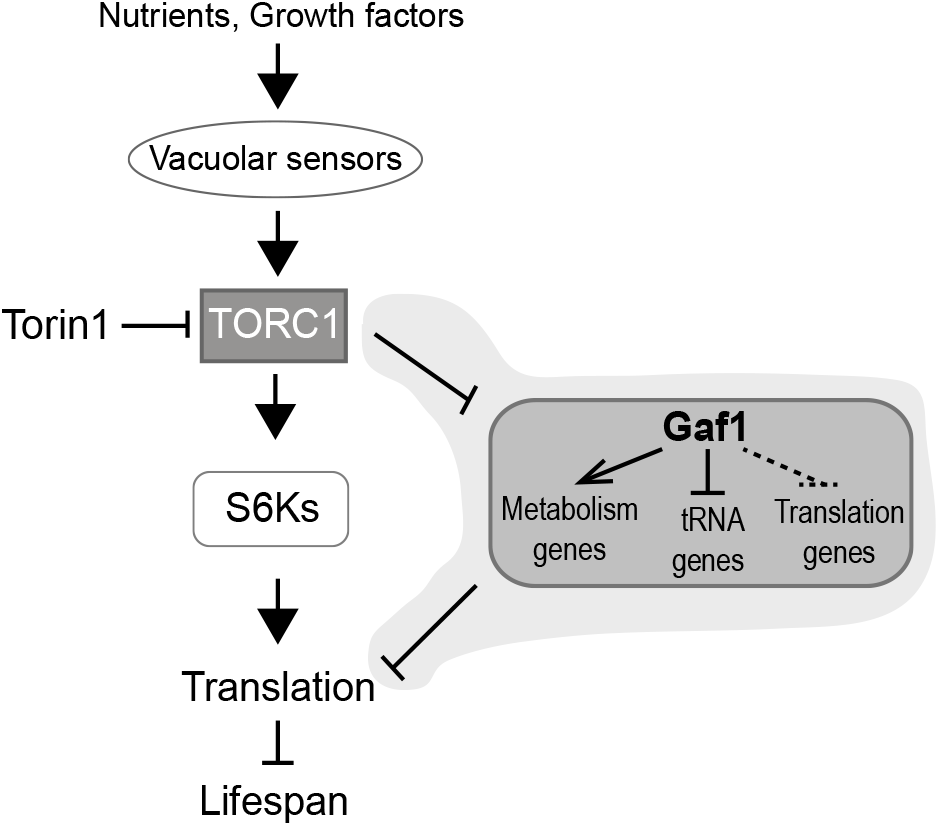
Model depicting transcriptional control of translation downstream of TORC1, mediated by Gaf1. Following TORC1 inhibition, Gaf1 activates the transcription of genes for small-molecule metabolic pathways and represses the transcription of tRNAs and other genes functioning in translation (the latter via indirect control, hatched). Together with the S6K-mediated translational control (Ma and Blenis, 2009), this transcriptional branch downstream of TORC1 contributes to longevity.

Inhibiting Pol III, which transcribes the tRNAs, prolongs lifespan in yeast, worms and flies, and is required for the lifespan extension mediated by TORC1 inhibition (Filer et al., 2017). Besides general transcription factors such as TFIIIB, TFIIIC and TBP, several factors control Pol III transcription without directly binding to DNA (Hummel et al., 2019), including the Pol III inhibitor Maf1, the coactivator PNRC, and MYC which interacts with the Pol III basal apparatus (Campbell and White, 2014; Graczyk et al., 2018; Zhou et al., 2007). To our knowledge, the TORC1 target Gaf1 is the first specific transcription factor shown to globally bind to and inhibit the tRNA genes. Thus, Gaf1 could mediate the ageing-associated function of Pol III. Recent work in flies shows that GATA transcription factors can mediate the effects of dietary restriction on lifespan (Dobson et al., 2018). This finding raises the possibility that Gaf1 regulation of ageing-related processes, such as tRNA transcription, is conserved, and that GATA factors exert similar functions downstream of TORC1 in other organisms. The mouse and human ortholog of Gaf1, GATA6, is involved in differentiation, stem-cell maintenance and cancer (Viger et al., 2008; Wamaitha et al., 2015; Zhong et al., 2011). It is plausible that GATA6 exerts these important functions by regulating translation-related genes, including tRNAs.

## Materials and Methods

### Strains and media

For wild-type control strains, we used *972 h^−^* or the parental strains for the deletion library, *ED666* (*h+ ade6-M210 ura4-D18 leu1–32*) and *ED668* (*h+ ade6-M216 ura4-D18 leu1–32*). All Bioneer haploid deletion library strains used were PCR-validated and backcrossed with *972 h^−^*. The *gaf1-GFP* strain was generated as described (Bahler et al., 1998). Cell cultures were grown in yeast extract plus supplements (YES) as default or in Edinburgh minimal medium (EMM2) if indicated (Moreno et al., 1991).Liquid cultures were grown at 32°C with shaking at 130 rotations per minute.

### Drug sensitivity assays

Cells were grown in liquid YES to an OD600 of 0.5. Ten-fold serial dilutions of cells were spotted onto YES agar plates, using replica platers for 48-well or 96-well plates (Sigma), with or without various drugs as indicated in figure legends.

### Measurement of cell size and fluorescence microscopy

To determine cell size, control and drug-treated cells were fixed in 4% formaldehyde for 10 min at room temperature, washed with 50 mM sodium citrate, 100 mM sodium phosphate, and stained with calcofluor (50 mg/ml). Microscopy was performed using a DAPI filter for calcofluor detection and a Hamamatsu ORCA-ER C4742-95 digital camera fitted to a Zeiss Axioskop microscope with EC plan-NEOFLUAR 63× 1.25 NA oil objective. Images were recorded using the Volocity acquisition program (PerkinElmer). At least 100 septated cells were counted and analyzed for each condition using the Volocity quantitation package (PerkinElmer). Results were analyzed using the R package. For fluorescence microscopy of Gaf1-GFP cells, we used a spinning disk confocal microscope (Yokogawa CSU-X1 head mounted on Olympus body; CoolSnap HQ2 camera [Roper Scientific], Plan Apochromat 100X, 1.4 NA objective [Olympus]). The images correspond to maximum intensity projections of 15 image stacks with a Z-step of 0.3 microns. Cells were immobilized with soybean lectin (Sigma L1395) in two different compartments of a glass-bottom 15 μ-Slide 8 well (Ibidi 80821) to add either DMSO as a solvent control or Torin1 (to a final concentration of 20 μM, dissolved in DMSO). In vivo chromatin staining was done with Hoechst 33342 (1 μg/ml). As this dye performs very poorly in YES, cells were immobilized onto glass bottom wells and washed three times with liquid EMM2 containing Hoechst 33342 (Sigma-Aldrich B2261) at 1 μg/ml plus Torin1 (20 μM). Cells were covered with this media and imaged 10 min later. Image analysis and editing was performed using Fiji (Image J) open software (Schindelin et al., 2012).

### Measurement of vacuolar size

Vacuolar labelling was performed as described (Codlin *et al*., 2009). Briefly, FM4-64 dye (Molecular Probes) was dissolved in DMSO at a concentration of 0.82 mM. Then, 2 μl FM4-64 stock was added to 1 ml log-phase cells with or without drugs. Following 30 min exposure to FM4-64, cells were washed, and chased for 40 min in fresh media to allow all dye to reach the vacuole. Fluorescence microscopy was performed using a Rhodamine filter for detection of FM4-64 and a Hamamatsu ORCA-ER C4742-95 digital camera fitted to a Zeiss Axioskop microscope with EC plan-NEOFLUAR ×63 1.25 NA oil objective. Images were recorded using the Volocity acquisition program (PerkinElmer). At least 500 vacuoles were measured using the Volocity quantitation package (PerkinElmer). Results were analysed using the R package.

### Chronological lifespan assay

Cells were grown in EMM2 media as described (Rallis et al., 2013). When cultures reached a stable maximal density, cells were left an additional 24 hrs and then harvested, serially diluted, and plated on YES plates. The measurement of colony-forming units (CFUs) was taken as timepoint 0 at the beginning of the lifespan curve (i.e., 100% cell survival). Measurements of CFUs were conducted on successive days until cultures dropped to 0.1% cell survival. Error bars represent standard deviation calculated from three independent cultures, with each culture measured three times at each timepoint. To determine the chronological lifespan when TOR is inhibited, 8 μM Torin1 was added to rapidly proliferating cell cultures at OD600 = 0.5 which were then grown to stationary phase, and lifespan was recorded as described above. AUCs were measured with ImageJ (Schindelin et al., 2012) for all experimental repeats using lifespans curves on the linear scale for % survival.

### High-throughput genetic screening

The haploid deletion libraries were plated onto YES plates containing 100 μg/ml G418 using a RoToR HDA robot (Singer). Multiple replicate copies of the library were thus generated. Using the RoToR, the libraries were compacted into nine 384-density plates of plates and then printed onto plates containing 20 μM Torin1. The plates were incubated at 32°C for 2 days and then manually scored for resistant colonies.

### Growth assay

Growth in the presence or absence of Torin1 were automatically determined in 48-well flowerplates using the Biolector microfermentation system (m2p-biolabs), at 1.5 ml volumes, 1000 rpm and 32°C. Growth dynamics were modelled using the grofit R package (Kahm et al., 2010). In the resulting growth curves, the units of the x-axis are time (hrs) while the y-axis shows biomass (arbitrary units) normalized to biomass at time 0.

### Western blotting and antibodies

For protein preparations, cells were diluted in 6 mM Na2HPO4, 4mM NaH2PO4.H2O, 1% Nonidet P-40, 150 mM NaCl, 2 mM EDTA, 50 mM NaF supplemented with protease (PMSF) and phosphatase inhibitors (Sigma cocktails 1 and 2), together with glass beads. Cells were lysed in a Fastprep-24 machine (MP Biomedicals). Phospho-(Ser/Thr) Akt Substrate (PAS) Antibody (9611, Cell Signaling) for detection of P-S6 (p27) and anti-rps6 (ab40820, Abcam) were used at 1/2000 dilution. For detection, we used the anti-rabbit HRP-conjugated antibody (1/5000 dilutions) with the ECL Western Blotting Detection System (GE Healthcare) according to the manufacturer’s protocol.

### Microarrays

Microarray analysis was performed as previously described (Rallis et al., 2013). Cells were grown in YES to OD600 = 0.5 and harvested. Torin1 treatments were done for 1 hr at a concentration of 20 μM. Caffeine/rapamycin treatments were also performed for 1 hr at concentrations 10mM Caffeine and 100ng/ml rapamycin. Two independent biological repeats with a dye swap were performed. For each repeat, a corresponding pool of Torin1 or caffeine/rapamycin treated and untreated wild-type and *gaf1Δ* cells was used as a common reference for microarray hybridization. Agilent 8× 15K custom-made *S. pombe* expression microarrays were used, with hybridizations and subsequent washes performed according to the manufacturer’s protocols. The microarrays were scanned and extracted using GenePix (Molecular Devices), processed using R scripts for quality control and normalization, and analyzed using GeneSpring GX3 (Agilent). We determined genes that were 1.5-fold up-regulated or down-regulated in both repeats of Torin1-treated and caffeine/rapamycin-treated *gaf1Δ* cells relative to Torin1-treated and caffeine/rapamycin-treated wild-type cells respectively.

### ChIP-seq

Cells were grown in YES to an OD600 of ~0.4. Untreated and Torin 1-treated (20 μM for 15 min or 1 hr) cells were fixed in 1% formaldehyde for 30 min and then quenched 10 min with 125mM glycine. Pellets were washed with ice-cold PBS, snap frozen in liquid nitrogen and stored at −80°C. Cell pellets were resuspended in lysis buffer (50 mM HEPES pH 7.6, 1mM EDTA pH 8, 150 mM NaCl, 1% Triton X-100, 0.1% sodium doxycolate, 1mM PMSF and protease inhibitors). Chromatin was obtained following cell disruption using a Fastprep-24 (MP Biomedicals) and sheared using a Bioruptor (Diagenode). Dynabeads M-280 sheep anti-rabbit IgG were incubated in lysis buffer and 0.5% BSA for 2 hrs with either rabbit anti-GFP (Abcam) for query IPs or 5 μl of rabbit-anti HA (Abcam) for control IPs. Then, 2 mg of Chromatin extract were inmunoprecipitated for 16 hrs using the corresponding antibody-incubated Dynabeads. Following the washes, DNA was eluted, treated with RNAse and proteinase K, and purified using the Qiagen PCR MiniElute kit. Sequencing libraries were prepared using the NEBNext^®^ ultra DNA Library Prep kit for Illumina^®^ (E7370L). DNA was sequenced using Illumina Mi-seq with a V3 kit, sequencing 75 bp on each end. Sequences were aligned to the *S. pombe* genome build ASM294v2 using Bowtie2. Peak calling was done with GEM (Guo et al., 2012) (setting --k_min 4 and --k_max 18), and peak annotation was done with the R package ChIPpeakAnno (Zhu et al., 2010). Peaks were annotated to the closest TSS; for peaks lying within 500 bp of 2 divergently expressed genes, peaks were annotated to both genes. Normalizations for the plots were performed using deeptools (Ramírez et al., 2016) (normalizing to RPGC and using the parameters –centerReads – binsize 10 –smoothLength 2). Further analyses were carried out with R scripts (http://www.r-project.org/). The ChIP-seq data are available from ENA at the following primary and secondary accession numbers: PRJEB32910 and ERP115647.

### Northern analyses

Detection of tRNA precursors was performed as described (Otsubo et al., 2018) using Digoxigenin labeled probes (Roche), following the manufacturer’s instructions. As a loading control, northerns were stripped by incubating for 60 min at 60°C with 0.1% SDS, changing the solution every 10 min, followed by re-hybridizing with a Digoxigenin labeled probe specific for *cdc2* (cdc2-SRT GGGCAGGGTCATAAACAAGC) as described (Clément-Ziza et al., 2014). Quantification of northern blots has been performed by ImageJ (Schindelin et al., 2012) as previously described (Rallis et al., 2014). Ratios of each tRNA band signal with the corresponding *cdc2* loading control have been normalized with the ratio at time point 0 for each tRNA and genotype.

## Acknowledgements

We thank Nazif Alic for critical reading of the manuscript, and Pawan Dhami (Genomics and Genome Engineering Facility funded by the Cancer Research UK-UCL Centre) for help with sequencing.

## Author Contributions

Conceptualization, C.R.; Methodology, C.R., S.G., M.R-L., V.A.T., S.C.; Investigation, C.R., M.R-L., S.G., O.H., V.A.T., E.T., S.C., J.B.; Formal Analysis, C.R., M.R-L., S.G., J.B. Writing, C.R., J.B.; Funding Acquisition, C.R., J.B.; Supervision: C.R., J.B.

## SUPPLEMENTAL MATERIAL

**Fig. S1.**
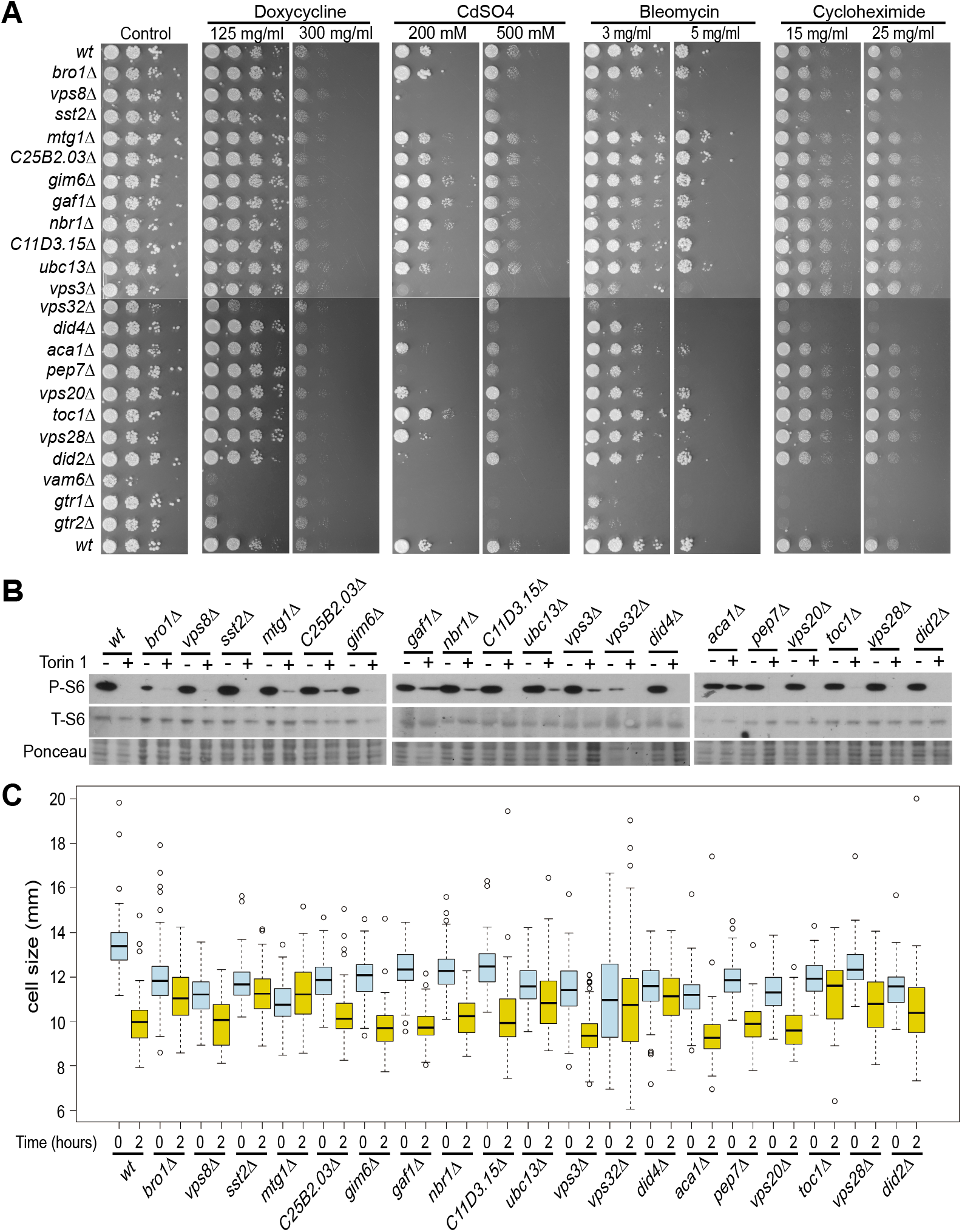
Characterization of Torin1-resistant strains. **A.** Resistance to Torin1 is not a result of multi-drug resistance. Spot assays of wild-type and the 19 resistant mutants in presence of different drugs as indicated. Control mutants that show multi-drug sensitivity are also spotted (*vam6Δ*, *gtr1Δ*, *gtr2Δ*) to show that the drugs are functional. **B.** Phosphorylation status of ribosomal S6 protein in verified Torin1-resistant mutants in the presence or absence of Torin1 as indicated. **C.** Cell size upon division of wild-type and resistant mutant cells, before (blue) and after (yellow) Torin1 treatment.

**Fig. S2.**
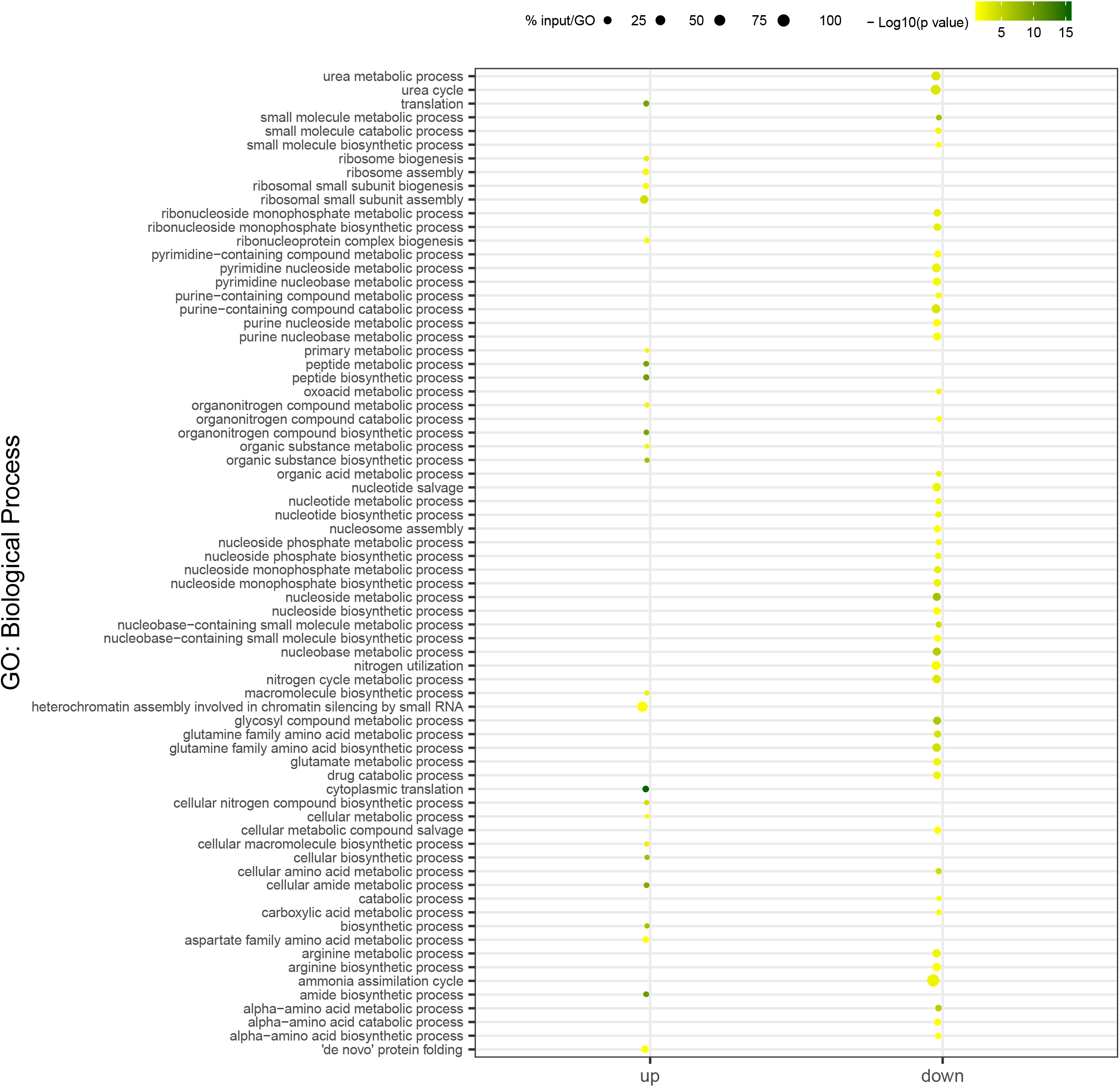
Visualization of all GO Biological Process categories enriched among the 108 genes that are repressed (down) and the 90 genes that are induced (up). Gene enrichment analysis was done using g:profiler (https://academic.oup.com/nar/advance-article/doi/10.1093/nar/gkz369/5486750). Results were plotted using a custom R script. Colors represent −log10 p values for the GO term enrichments. The size of the bubbles represents the ratio between the number of genes in the gene list and the total number of genes in the GO category (in percentage); only terms with p values <0.05 are represented.

**Fig. S3.**
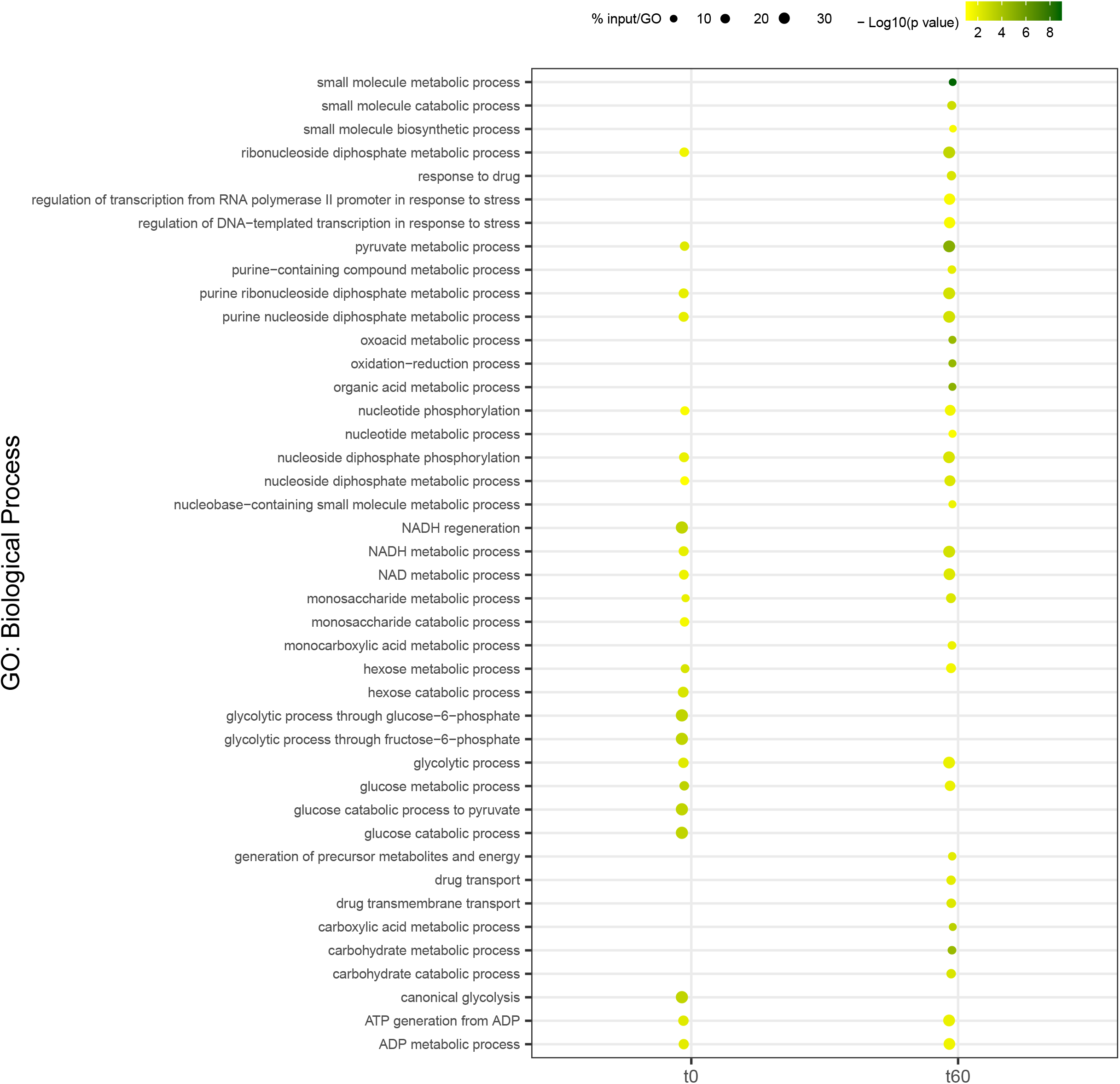
Visualization of all GO Biological Process categories enriched among the 245 protein-coding genes whose promoters are bound by Gaf1 after 60 min of Torin1 treatment. See Fig. S2 for details.

**Fig. S4.**
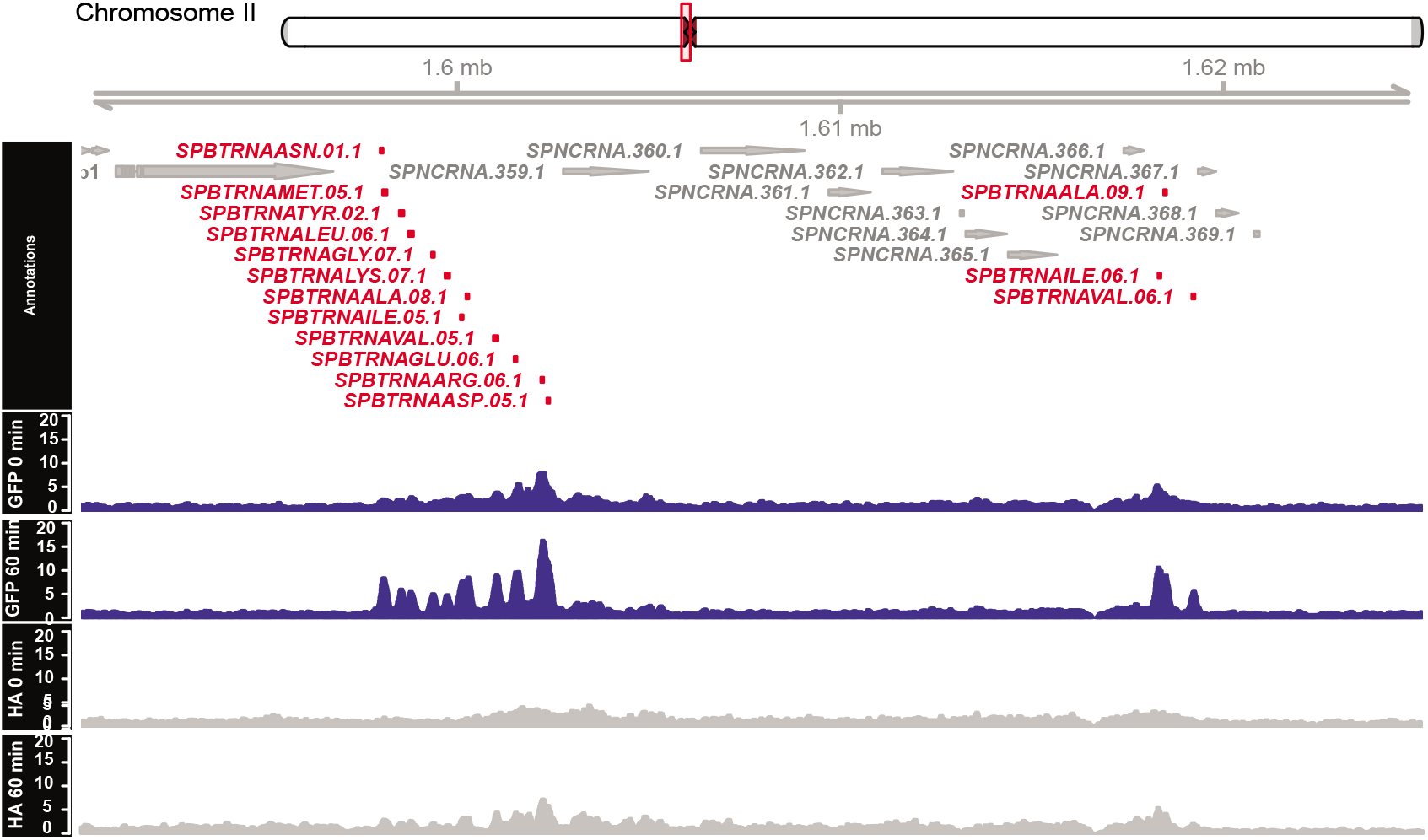
Gaf1 binding peaks within Chromosome II region containing clustered tRNA genes (marked in red). Binding profiles are shown for cells before (0 min) and 60 min after treatment with Torin 1 as indicated. Signals from experimental IPs (GFP) are shown in blue, while control IPs (HA) are shown in grey.

**Table S1.** List of the 19 mutants that show resistance to Torin1. Mutants with black fonts are shown previously to be resistant to Torin1. Mutants with red fonts are first reported as Torin1-resistant in this study.

**Table S2.** Lists of genes that are ≥1.5 fold-differentially expressed in *gaf1Δ* compared to wild-type cells following 60 min treatment with 20 μM Torin1

**Table S3.** List of genes whose promoters are bound by Gaf1 before and 60 min after Torin1 treatment.

